# A type II secreted subtilase from commensal rhizobacteria disarms the immune elicitor peptide flg22

**DOI:** 10.1101/2024.05.07.592856

**Authors:** Samuel Eastman, Ting Jiang, Kaeli Ficco, Chao Liao, Britley Jones, Sarina Wen, Yvette Olivas Biddle, Aya Eyceoz, Ilya Yatsishin, Todd A. Naumann, Jonathan M. Conway

## Abstract

Plant roots grow in association with a community of microorganisms collectively known as the rhizosphere microbiome. Immune activation in response to elicitors like the flagellin-derived epitope flg22 restricts bacteria on plant roots but also inhibits plant growth. Some commensal root-associated bacteria are capable of suppressing the plant immune response to elicitors. In this study, we investigated the ability of 165 root-associated bacteria to suppress flg22-induced immune activation and growth restriction. We demonstrate that a type II secreted subtilase, which we term Immunosuppressive Subtilase A (IssA), from *Dyella japonica* strain MF79 cleaves the immune eliciting peptide flg22 and contributes to immune suppression. IssA homologs are found in other plant-associated commensals, with particularly high conservation in the order Xanthomonadales. This represents a novel mechanism by which commensal microbes modulate flg22-induced immunity in the rhizosphere microbiome.

## INTRODUCTION

Plant roots are colonized by a microbiome of pathogenic, mutualistic, and commensal bacteria which collectively influence the growth and development of the host ^1^. Intracommunity interactions between rhizobacteria strongly shape community composition and host phenotypes ^1,2^. Commensal bacteria comprise the bulk of the rhizosphere microbiome and can be beneficial or detrimental to the host plant depending on genetic and environmental factors that are not fully understood ^3,4^.

Rhizobacteria – pathogenic, mutualistic, or commensal – must contend with the plant immune system once they transition from soil to the plant root ^5-7^. Bacteria possess conserved microbe-associated molecular patterns (MAMPs) which are detected by plant pattern recognition receptors (PRRs): a process that elicits a basal immune response termed MAMP-triggered immunity (MTI) ^8^. Flg22, a potent and well-studied MAMP, is a 22-residue polypeptide derived from bacterial flagellin by plant proteases that stimulates MTI when bound by the flg22 receptor FLS2 (FLAGELLIN-SENSING 2) and co-receptor BAK1 (BRI1-ASSOCIATED RECEPTOR KINASE 1) ^9-11^.

The 17 amino acids at the N-terminus of flg22 constitute an “address” that interacts with FLS2 while the five amino acids at the C-terminus of flg22 constitute a “message” that is jointly bound by FLS2 and BAK1 and drives receptor complex formation and MTI activation ^12-14^. MTI encompasses a broad set of plant immune responses including the production of reactive oxygen species (ROS) and phytoalexins like camalexin which nonspecifically kill or restrict microbes ^8,15,16^. Plant immunity is resource-intensive and, thus, defense activation occurs at the expense of growth ^17-19^. For instance, prolonged exposure to flg22 inhibits root growth, a phenomenon known as root growth inhibition (RGI) ^20^. Although it is still unclear how plants can cooperate with beneficial microbes while simultaneously restricting pathogens, recent evidence has indicated that some commensal bacteria suppress plant immunity, contributing to a proper balance of growth and defense ^6,21,22^. One of the primary microbial strategies to suppress plant immunity is the secretion of effectors into host cells or the apoplast ^23^. In bacteria, apoplastic effectors are often secreted by the type II secretion system (T2SS) ^24^. Phytobacterial type II effectors are typically degradative enzymes like cellulases, pectate lyases, polygalacturonases, and xylanases that break down the cell wall ^25^. A few plant pathogens are known to utilize type II apoplastic serine proteases, but the functions of these serine proteases are not clear ^26-28^. ROS produced as a plant MTI response attenuates type II secretion and “tames” a conditionally pathogenic *Xanthomonas* sp. by reducing virulence, suggesting that type II secreted effectors are important for the virulence of phytobacteria in order Xanthomonadales ^29^. While pathogenic, commensal, and mutualistic bacteria all secrete effectors, the mechanisms by which commensal bacteria regulate plant immunity are largely unknown ^30-32^.

Serine proteases are a broad class of proteases distinguished by their use of serine as a catalytic residue ^33^. One of the largest families of serine proteases is the S8 (or subtilase) family, which is further divided into the S8A (subtilisin) and S8B (kexin) subfamilies ^34^. Almost all subtilases are endopeptidases which utilize an Asp-His-Ser (DHS) catalytic triad ^33^. Subtilases are abundant in plants and important to crucial functions, including the immune response ^35^. A number of plant subtilases positively activate plant immune signaling cascades or plant cell death in response to pathogens ^35^. In contrast, the *Arabidopsis thaliana* site-1 protease (S1P) cleaves the propeptide of the endogenous rapid alkalinization factor 23 (RALF23), a process which inhibits PAMP receptor complex formation and thus negatively regulates immunity ^36,37^. Some bacterial and fungal subtilases act as PAMPs and activate plant immunity independent of their enzymatic activity ^38,39^. However, no bacterial subtilases are known to negatively regulate plant immunity.

Order Xanthomonadales (also called Lysobacterales) within phylum Gammaproteobacteria is comprised of diverse bacteria from many environments, including many plant pathogens and commensals ^29,40^. Within order Xanthomonadales, the genus *Dyella* and type species *Dyella japonica (Dja)* were originally described as soil isolates ^41^. *Dja* strain MF79, which was isolated from surface-sterilized Arabidopsis roots, is a robust endophytic colonizer of Arabidopsis roots in synthetic community studies ^42-44^. *Dja* MF79 strongly suppresses plant MTI and promotes colonization by other bacteria in a type II secretion-dependent manner ^21^. *Dja* MF79 culture supernatant retained by, but not passed through, a 10 kDa filter is capable of suppressing flg22-induction of a *pCYP71A12::GUS* reporter in Arabidopsis, suggesting the presence of a secreted type II effector capable of interfering with the plant immune response to flg22 ^21^. In this study, we screened a collection of 165 root-associated bacterial isolates and defined strong and weak suppressors of flg22-induced plant immunity. We identify a type II-secreted effector subtilase which rapidly cleaves the immune elicitor peptide flg22 and is partially responsible for strong suppression of flg22-induced immunity by *Dja* MF79 and other commensal rhizobacteria with suppressive phenotypes.

## RESULTS

### Root-associated bacteria differentially suppress flg22-induced root growth inhibition and *pCYP71A12* activation

To assess the ability of root-associated bacteria to suppress plant immunity, we utilized a collection of 165 bacterial isolates from Brassicaceae roots (primarily Arabidopsis) grown in two natural soils from North Carolina ^45^. The collection is comprised of 33 bacterial genera belonging to the 4 dominant phyla of the plant rhizosphere microbiome (Table S1). We screened the collection for suppression of flg22-induced root growth inhibition (RGI) of an *Arabidopsis thaliana* line (*pWER::FLS2-GFP*) which strongly expresses FLS2 in root tissues ^46^, a well-established method for assessing the effects of root-associated bacteria on plant immune responses ^20-22^. Seven-day-old germ-free Arabidopsis seedlings were co-cultured with individual bacterial isolates for an additional seven days on 0.5× MS agar plates containing 1 μM flg22. Arabidopsis seedlings co-cultured with 41% (67 out of 165) of isolates in the collection exhibited significantly greater root elongation than uninoculated controls (*P* value <0.5, T statistic <0), indicating that taxonomically diverse bacteria can suppress flg22-induced RGI (Figure 1A-B, Table S1-3).

**Figure 1:**
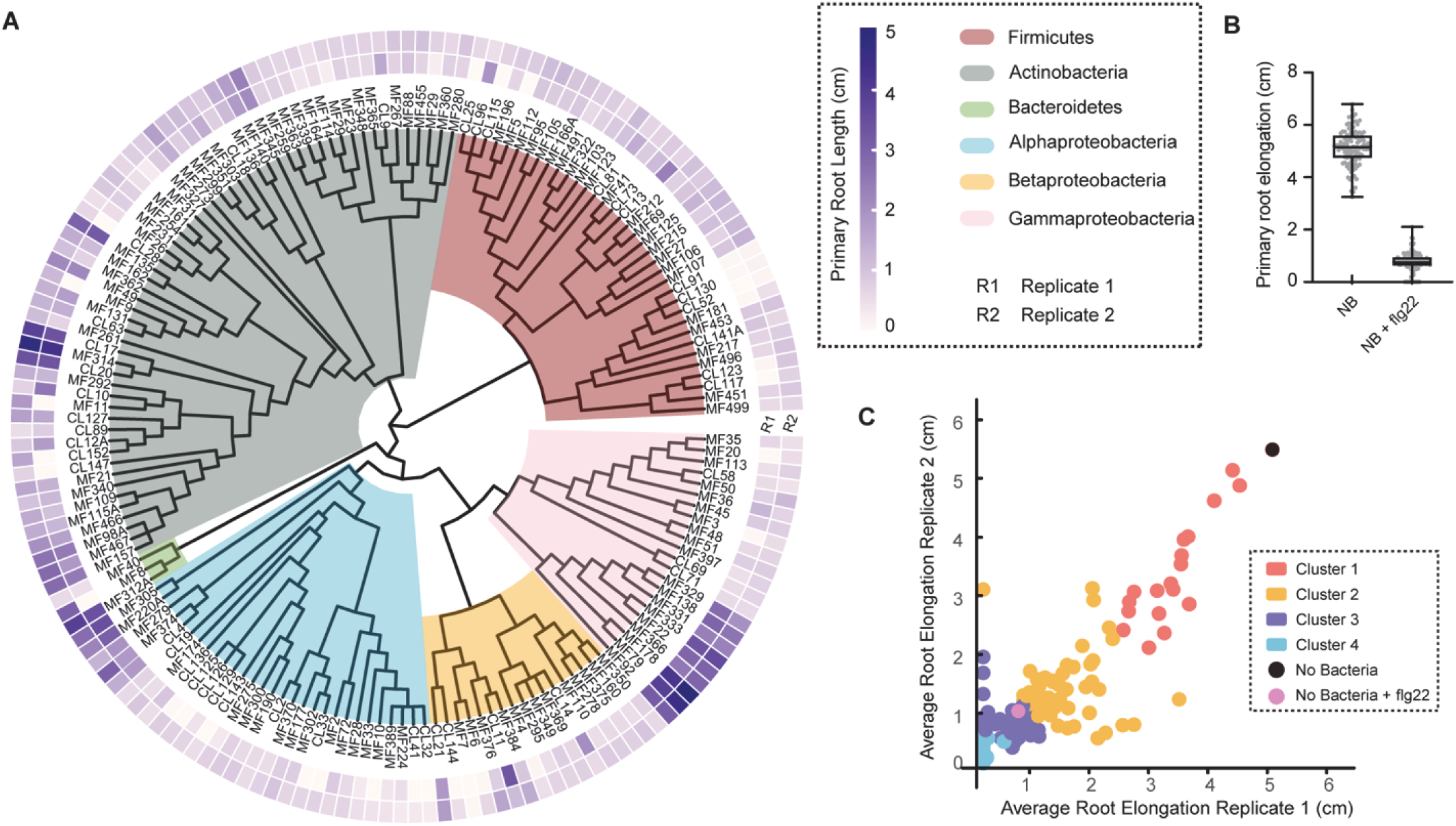
Identification of strains with RGI suppressive activity from 165 root-associated bacteria isolates. (A) Phylogenetic tree of 165 root-associated bacterial strains isolated from Brassicaceae roots, primarily *Arabidopsis*. Two heatmap rings represent two biological replicates, illustrating the average primary root elongation of *Arabidopsis* line *pWER::FLS2-GFP* when cocultured with individual strains on 0.5xMS plates with 1 μM flg22. The middle text ring indicates strain IDs, while the inner ring shows phylogenetic relationships colored by phylum or class (Proteobacteria) levels. (B) Primary root elongation of seedlings on 0.5x MS plates with or without flg22 (n = 105). ‘NB’ indicates plates without bacteria. (C) Scatter plot of all 165 strains and no bacterial controls based on their two replicates of average root elongation. Strains with similar RGI suppressive activity are colored according to K-means clustering analysis. The optimal number of clusters was determined using the elbow method.

We employed K-means clustering to categorize isolates based on their RGI suppressive activity, determining the optimal number of clusters using the elbow method. We classified all 165 isolates into four distinct clusters (Figure 1C), with 67 isolates exhibiting significant suppressive function grouped within Cluster 1 and Cluster 2 (Table S1). The 18 isolates comprising Cluster 1 displayed robust suppressive activity, with an average root length of 3.21 cm (Table S1). Taxonomically, these isolates show specific distribution at the order level, with six isolates belonging to Micrococcales and nine to Xanthomonadales. Among our 165-member collection, not all isolates from Micrococcales are categorized in Cluster 1 but, intriguingly, all isolates from Xanthomonadales exhibit strong suppressive activity and are grouped in Cluster 1. This includes all members in the collection from the genera *Stenotrophomonas*, *Luteibacter*, and *Dyella*. Our results are consistent with a recent study ^22^ that screened a distinct microbial isolate collection for the ability to suppress flg22-induced RGI and similarly found that all tested isolates from Xanthomonadales, including the genera *Lysobacter*, *Pseudoxanthomonas*, and *Rhodanobacter*, acted as RGI suppressors (Tables S1 and S4). These findings suggest that a conserved mechanism for flg22-induced RGI suppression is present in bacteria in order Xanthomonadales.

We next assessed the ability of 25 of the strongest suppressors of flg22-induced RGI to suppress flg22-induced *pCYP71A12::GUS* activation, another commonly used method for assessing plant immune modulation by commensal bacteria ^21,47^. Six day-old Arabidopsis *pCYP71A12::GUS* seedlings, which express the GUS reporter gene specifically in the root elongation zone in response to flg22, were inoculated with bacteria at densities ranging from OD_600_=2×10^-1^ to 2×10^-6^, co-cultured in liquid 0.5× MS media overnight, and treated with 100 nM flg22 (Figure S1). The majority of tested isolates suppressed flg22-induced GUS expression, and some isolates were able to suppress GUS expression at bacterial densities as low as OD_600_=2×10^-5^ (Figure 2B). Surprisingly, the ability of each isolate to suppress flg22-induced RGI does not appear to be strongly correlated with the ability to suppress flg22-induced *pCYP71A12::GUS* activation (Figure 2). *Brevundimonas* sp. MF374, in Cluster 2, suppressed flg22-induced *pCYP71A12::GUS* activation with an inoculation density of OD_600_=2×10^-5^. In contrast, the three strongest RGI suppressors overall, the Cluster 1 isolates *Curtobacterium* sp. MF314 and the near-isogenic *Dyella japonica* isolates MF178 and MF79, suppressed flg22-induced *pCYP71A12::GUS* activation only at bacterial densities as low as OD_600_=2×10^-3^ and 2×10^-4^, respectively (Figure 2, Figure S1).

**Figure 2:**
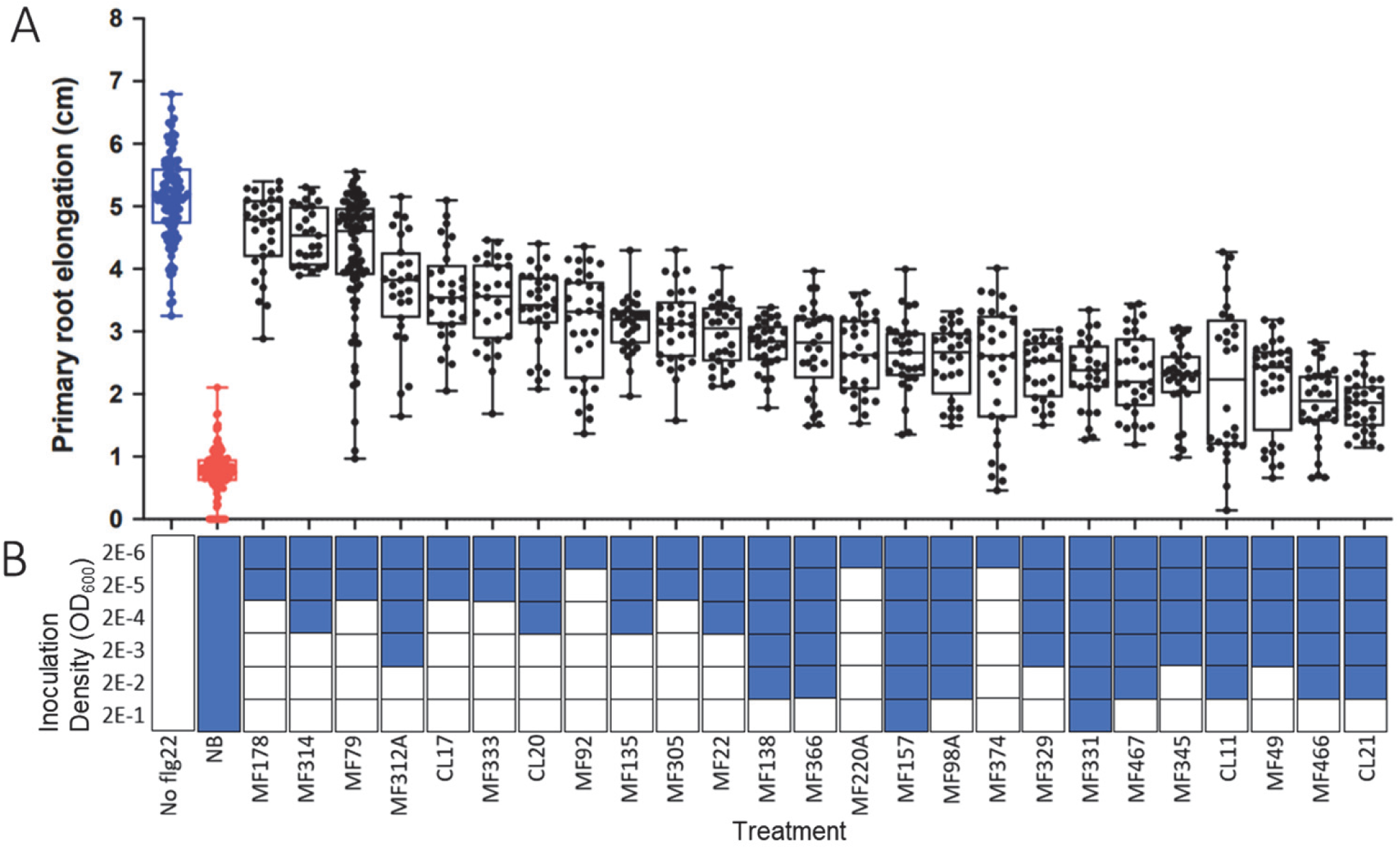
Commensal root microbes differentially suppress aspects of flg22-induced immunity. (A) Root growth inhibition (RGI) assay. Arabidopsis *pWER:FLS2-GFP* plants were germinated on sterile growth media plates for one week then transferred to plates containing 1 µM flg22 spread with 100 µL bacteria (OD_600_=0.001) and the length of their primary root was marked. The plants were grown for one additional week then the elongation of the primary root was measured. (B) GUS assay. Arabidopsis *pCYP71A12*::*GUS* plants were germinated in sterile growth liquid media for six days, then the growth media was replaced with media containing bacteria at the indicated densities. The following day, plants were induced with 100 nM flg22 for five hours then incubated with GUS reagent overnight. Plants were destained and root tips were photographed under a microscope. A blue box indicates the majority of roots displayed a flg22-induced *pCYP71A12* pattern of *GUS* activation while a white box indicates the majority of roots displayed a naïve phenotype. (See also Figure S1)

### *Dja* MF79 contains a putative subtilase with homology to Xcv3671

In addition to suppressing flg22-induced RGI in our screens, *Dja* MF79 has been shown to suppress flg22-induced *pCYP71A12::GUS* activation in a type II secretion-dependent manner ^21^. To identify potential type II secreted effectors (T2SEs) in *Dja* MF79, we considered other bacteria in order Xanthomonadales that had been shown to suppress plant immunity via T2S substrates. *Xanthomonas campestris* pv. *vesicatoria* (*Xcv*) secretes T2SEs – including a lipase, two xylanases, and a protease – that contribute to *Xcv* virulence on host pepper and tomato ^28,48^. We utilized BLASTp ^49^ to search the *Dja* MF79 genome for proteins homologous to effectors from *Xcv* strain 85-10 ^28,48^. *Xcv* T2S protease Xcv3671 is homologous to a putative subtilase in *Dja* MF79 encoded by IMG ^50^ GeneID 2558297680, particularly around the putative catalytic DHS triad characteristic of subtilases ^28,51^. The subtilase, which we term Immunosuppressive Subtilase A (IssA) is predicted to have a cleaved Sec/SPI signal peptide by SignalP 6.0, consistent with putative type II secretion ^52^. The *issA* gene is encoded 5 kb upstream of the T2SS gene cluster of *Dja* MF79 which, when disrupted by a transposon insertion in either *gspE* or *gspD* (IMG GeneID 2558297684 and 2558297694, respectively), strongly reduces suppression of flg22-induced immunity ^21^. This could indicate that *issA* is expressed in tandem with type T2SS machinery.

### IssA suppresses flg22-induced *pCYP71A12::GUS* activation

To assess the contribution of IssA to *Dja* MF79 immune suppression, we generated *Dja* MF79 Δ*issA* mutants and conducted RGI and GUS assays. The Δ*issA* knockout mutant and a previously described general secretory pathway D transposon insertion mutant (*gspD*::Tn) ^21^ failed to suppress 100 nM flg22-induced *pCYP71A12:GUS* activation at bacterial inoculation densities of OD_600_=2×10^-3^ (Figure 3A, Figure S2A). Wild type *issA* expressed on a replicating plasmid complemented the Δ*issA* mutant phenotype, but *issA* with alanine substitutions in the conserved DHS catalytic triad did not (Figure 3A). Surprisingly, suppression of flg22-induced RGI by *Dja* MF79 was seemingly unaffected by *issA* deletion (Figure S2B).

**Figure 3:**
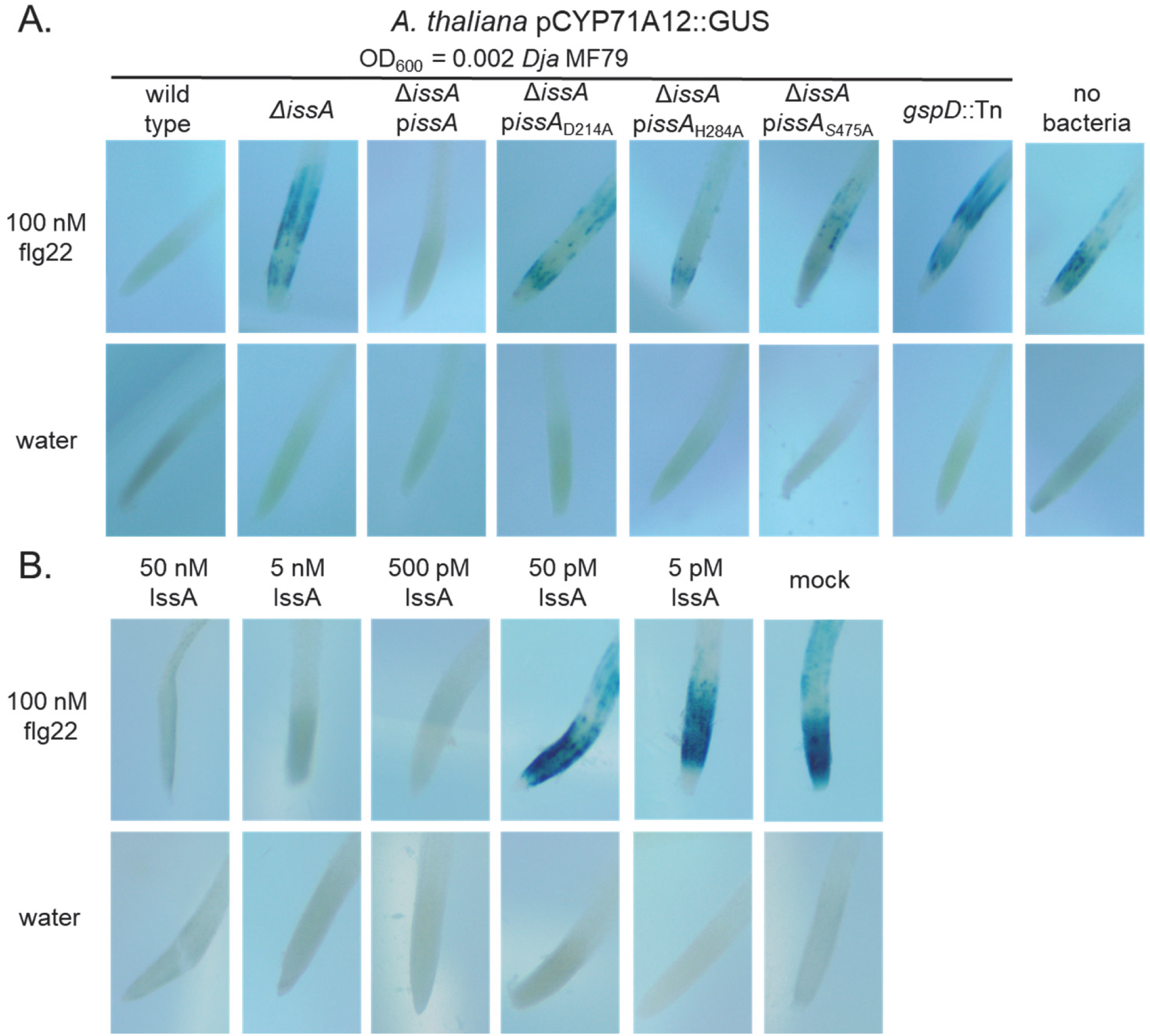
IssA suppresses flg22-inducted immunity. (A) GUS assay utilizing Arabidopsis *pCYP71A12*::*GUS* roots and *Dyella japonica* strains. Wild-type and *issA-*complemented *Dja* suppress induction of *pCYP71A12*::*GUS* by 100 nM flg22, but Δ*issA* and catalytic mutants do not. Plants were grown aseptically for six days, grown with bacteria for one day, co-incubated with flg22 for five hours, and incubated with GUS reagent overnight. (B) GUS assay utilizing purified, recombinant IssA. IssA suppresses *pCYP71A12*::*GUS* induction by 100 nM flg22 at concentrations as low as 500 pM. Plants were grown aseptically for six days, co-incubated with flg22 and IssA for 5 hours, and incubated with GUS reagent overnight.

To directly test the ability of IssA to suppress flg22-induced *pCYP71A12::GUS* activation, we expressed and purified recombinant IssA from the methylotrophic yeast *Pichia pastoris* (syn. *Komagataella phaffii*). We observed recombinant IssA as a doublet when analyzed by SDS-PAGE (Figure S3A). The bands were smaller than the expected 52 kDa, and LC-MS/MS analysis of their tryptic peptides identified N-terminal tryptic peptides beginning with T157 (Figure S3B). This is consistent with two structurally homologous bacterial subtilases from the protein data bank ^53,54^. Limited degradation of IssA occurred during purification, as reported previously for other subtilases ^55-57^. In bioassays, purified, recombinant IssA suppressed *pCYP71A12::GUS* activation by 100 nM flg22 at concentrations of 500 pM or greater (Figure 3B). Our data indicates that IssA is an enzymatically active subtilase that inhibits flg22-induced immune responses.

### IssA homologs are distributed in other immune suppressive root-associated bacteria

IssA-like subtilases are not confined to genus *Dyella*, and homologs can be identified in a number of other plant-associated bacteria. We performed tBLASTn searches of IssA against the genomes of the 165-member isolate collection. We found 398 proteins from 110 bacterial genomes with homology to IssA greater than the E-value cutoff of 1.00E-5, with the majority of these proteins being annotated as serine proteases or subtilases (Table S5). Of these, 313 proteins from 101 genomes were predicted to contain a signal peptide by SignalP6.0 (Table S5) ^52^. The closest IssA homologs were found in other immunosuppressive Gammaproteobacteria, including *Luteibacter* spp., *Stenotrophomonas* spp., and other *Dyella* spp., however high-scoring homologs were also found in the Actinobacteria, including *Arthrobacter* spp. and *Paenarthrobacter* spp., and in the Betaproteobacteria, including *Burkholderia* spp. and *Ralstonia* spp. (Figure 4A, Table S5). Of the 22 bacterial isolates capable of limiting flg22-induced RGI to 1.6 cm or greater, only five lacked a secreted IssA homolog – most notably three isolates of *Curtobacterium* spp. (Figure 4A, table S5).

**Figure 4:**
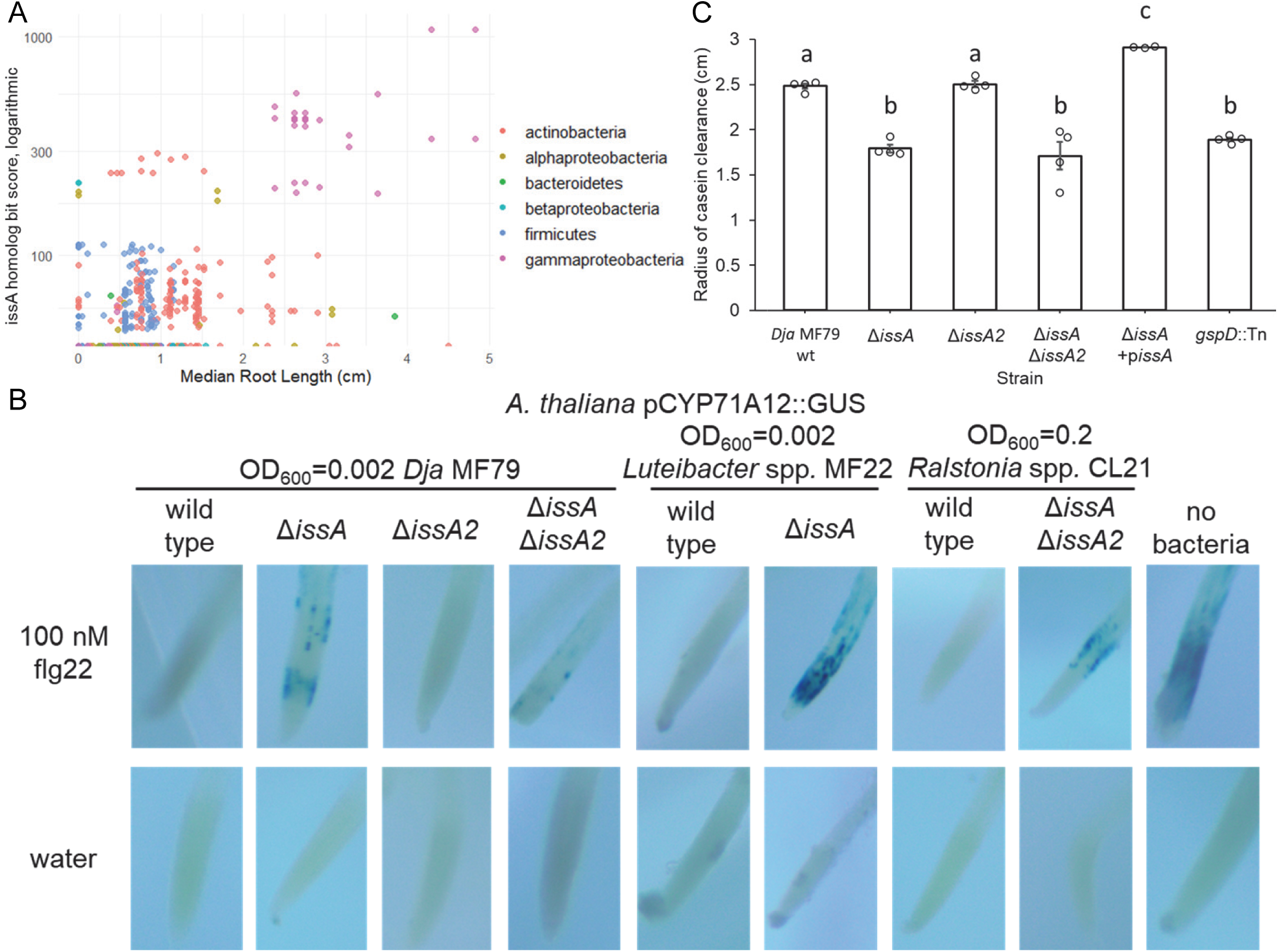
IssA, but not IssA2, homologs are active subtilases which suppress flg22-induced immune responses. (A) Scatterplot graphing the similarity of IssA homologs (in bits) against the root lengths of Arabidopsis *pWER:FLS2-GFP* seedlings exposed to 1 µM flg22 and bacteria strains at OD_600_=.001 (see Table S2, Table S5). Homologs were found by BLAST searching the genomes of 165 bacterial isolates and filtering for proteins with predicted secretion using SignalP6.0. A bit score of 0 indicates the strain contains no homologs. Homologs are colored by taxonomy. (B) GUS assays of bacterial strains with deletions in *issA* and/or *issA2*. *pCYP71A12::GUS* seedlings were inoculated with the minimum bacteria density required for the wild-type strain to (Figure 2). *issA,* but not *issA2,* suppression causes a reduction in suppression of flg22-induced *pCYP71A12:GUS* activation. (C) Radius of casein clearance by *Dja* MF79 strains. Deletion of *issA*, but not *issA2*, significantly reduced the clearance halo to a similar level as a *gspD* transposon insertion mutant. Complementation with plasmid-expressed *issA* significantly increased casein clearance beyond the wild-type. This experiment was repeated twice with similar results. Bars indicate standard error. Letters indicate statistical significance of p<.05 in an ANOVA with a Tukey post-hoc test.

*Dja* MF79 and closely related bacteria also contain among the homologs a second putative secreted subtilase, which we tentatively labeled IssA2, which is encoded adjacent to IssA in the genomes of some bacteria like *Ralstonia* sp. CL21 (Table S5). We constructed Δ*issA* and/or Δ*issA2* mutants in suppressive isolates *Dja* MF79 (*issA2*, IMG Gene ID 2558295051), *Luteibacter* sp. MF22 (*issA*, IMG Gene ID 2520562849), and *Ralstonia* sp. CL21 (*issA/issA2*, IMG Gene IDs 2558854277 and 2558854276). Δ*issA* mutants demonstrated a roughly 10-fold increase in bacterial density needed to suppress flg22-induced *pCYP71A12::GUS* activation at the same level as the wild type strain, while Δ*issA2* mutants showed no change in suppression from the wild type (Figure 4B, Figure S4). *Dja* MF79 Δ*issA*, but not Δ*issA2*, mutants also showed a reduced clearance radius on LB agar plates supplemented with 1% casein, suggesting that IssA, but not IssA2, has general protease activity (Figure 4C). Our data indicates that IssA homologs in *Luteibacter* and *Ralstonia* spp. also act as suppressors of flg22-induced plant immunity, and that suppression of flg22-induced plant immunity is not a trait shared by all secreted subtilases.

### IssA alters flg22 immunogenicity *in vitro*

*Dja* MF79 cell-free supernatant retained by, but not passing through, a 10 kDa Molecular Weight Cut-Off (MWCO) filter was previously shown to suppress flg22-induced *pCYP71A12* activation, indicating the presence of a secreted protein effector ^21^. We replicated this experiment and found the *Dja* MF79 cell-free supernatant suppressed the response to flg22 (Figure S5A-C). We then co-incubated *Dja* MF79 cell-free supernatant with flg22 for one minute before passing the mixture through a 10 kDa MWCO filter to remove IssA and recover flg22 or altered flg22 products. We then applied the filtrate to *pCYP71A12::GUS* plants at an estimated concentration of 100 nM flg22 and found that flg22 co-incubated with wild type or *issA*-complemented *Dja* MF79 culture supernatant failed to induce *pCYP71A12::GUS*, while flg22 co-incubated with Δ*issA* or *gspD::Tn* culture supernatant, as well as with uninoculated culture supernatant, still induced GUS expression (Figure S5B-C). Our data indicates IssA alters the immunogenicity of flg22 and that direct interaction of IssA with plant cells is unnecessary for immune suppression.

We co-incubated flg22 with purified, recombinant IssA and analyzed reaction time points by MALDI-TOF mass spectrometry. We found that IssA initially cleaved flg22 between either N10-S11, resulting in QRLSTGSRIN and SAKDDAAGLQIA peptides, or D14-D15, resulting in QRLSTGSRINSAKD and DAAGLQIA peptides, each observed as multiple molecular ion adducts (Figure 5). Extended incubation led to subsequent cleavage at two secondary sites, S7-R8 and A17-G18, observed as loss of larger peptides and generation of peptides RIN and GLQIA and QRLSTGS (Figure 5). We did not observe peptide ions that would indicate cleavage at any additional peptide bonds. Together, our data indicates that IssA cleaves flg22 *in vitro* to produce specific product peptides, altering its immunogenicity and precluding plant immune activation.

**Figure 5:**
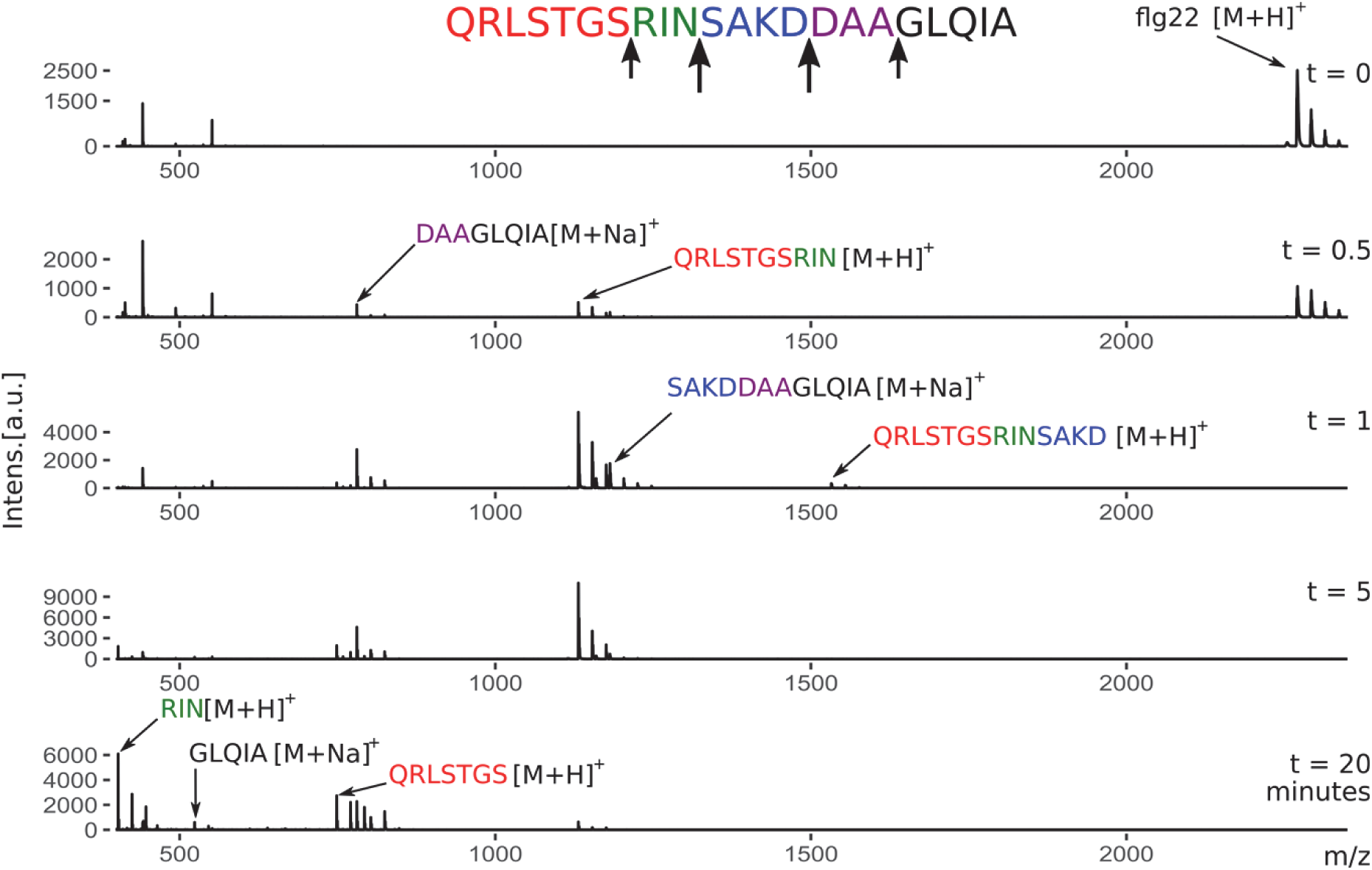
IssA cleaves flg22 at several sites *in vitro*. MALDI-TOF MS chromatograms for mixtures of flg22 and IssA. Substrate peptide flg22 (1 mM) was incubated with purified recombinant IssA (1 µM) and reaction products were visualized at different time points. Product peptides corresponding to different cleavage sites of flg22 by IssA are indicated in different colors.

## DISCUSSION

In this study, we used a simple plant-microbe interaction system to screen a collection of 165 bacterial isolates isolated from plant roots for stains capable of reversing the flg22-induced RGI. Of these isolates, 41% (67 out of 165) exhibited significantly longer roots compared to the non-bacterial control (Figure 1). The taxonomic diversity of these 67 suppressive isolates suggests that many bacteria are capable of suppressing the plant immune system. Through clustering analysis, we identified 18 of 67 isolates as strong suppressors, with an average root elongation of 3.21 cm in the RGI assay. These isolates are not highly phylogenetically diverse but are concentrated in the orders Micrococcales (six isolates) and Xanthomonadales (nine isolates), which are common and abundant in the rhizosphere microbiome. All of the order Xanthomonadales isolates we screened were strong suppressors, aligning with a similar study of a distinct collection of Arabidopsis microbiome isolates in which all Xanthomonadales screened were also suppressors (Table S4) ^22^.

We report that the type II-secreted subtilase IssA contributes to suppression of flg22-induced plant immunity by root-associated bacterium including *Dyella japonica* MF79. A homology search of the MEROPS protein database (https://www.ebi.ac.uk/merops/) finds that IssA and its *Xanthomonas* homolog Xcv3671 most closely resemble subtilases containing an MprA domain (data not shown), a group which include proteases with broad substrate specificity including the proteins casein, keratin and collagen ^58-60^. Xcv3671 is thought to have narrow substrate specificity, as deletion of Xcv3671 does not significantly change the ability of *Xcv* to digest casein ^28^, but the exact substrate of Xcv3671 that enables it to suppress plant immunity is unknown. In contrast, we found that *issA* deletion significantly reduced casein cleavage by *Dja* MF79 to levels similar to the *gspD*::Tn mutant (Figure 4C), indicating IssA has a broader substrate specificity than the flg22 peptide and may digest other peptides found at the root-microbe interface.

*Dja* MF79 likely secretes at least one additional suppressive protein in addition to IssA, as the effects of *issA* deletion on suppression of flg22-induced *pCYP71A12*::*GUS* were less than transposon insertion in *gspD* (Figure S2A) ^21^. In the genome wide transposon mutagenesis screen preformed previously ^21^, only transposon insertion in the T2SS (*gspD*::Tn and *gspE*::Tn) resulted in the loss of suppression of *pCYP71A12::GUS* activation, suggesting complementary suppressive molecules are secreted by *Dja* MF79. We show IssA is one such secreted component of flg22-activated immune suppression. Deletion of the homologous *issA2* gene had no effect alone or in combination with *issA* deletion, indicating the additional secreted suppressive molecules remain to be discovered.

While we observed a significant reduction in the ability of bacterial *ΔissA* mutants to suppress flg22-induced *pCYP71A12::GUS* activation, we did not observe a corresponding reduction in suppression of flg22-induced RGI (Figure 3, Figure S2). This further supports the conclusion that additional suppressive molecules may be secreted by *Dja* MF79. More broadly, we observed that bacterial suppression of flg22-induced RGI was not strongly correlated with suppression of flg22-induced *pCYP71A12::GUS* induction (Figure 2). This suggests that these two commonly used assays are not directly comparable, and that the dynamics of bacterial suppression of flg22-induced immune activation may be affected by other variables such as duration of interaction and whether the root is submerged in an aqueous environment. In our study, we tested the ability of bacterial root isolates to suppress flg22-induced *pCYP71A12::GUS* activation at inoculation densities of OD_600_=2×10^-1^ to 2×10^-6^ (Figure 2B, Figure S1). Our method represents an improvement in sensitivity over previously performed assays ^21,47^, which only assessed the ability of bacteria to suppress flg22-induced immune activation at an inoculation density of OD_600_=2×10^-3^.

We were unable to obtain soluble purified IssA using conventional bacterial protein expression in *E. coli* (data not shown). To produce purified proteins for testing in bioassays (Figure 3B) and *in vitro* protease assays (Figure 5) we instead heterologously expressed IssA in the methylotrophic yeast *P. pastoris*. *P. pastoris* is commonly used to produce secreted eukaryotic proteins including those from plants and fungi. Our work demonstrates that, despite being a eukaryote, *P. pastoris* can be a useful host for producing type II secreted bacterial proteins.

We observed that purified, recombinant IssA cleaved flg22 at four different sites at varying rates (Figure 5). These cleavage events separate the N-terminal 17 amino acid “address” of the flg22 peptide from the C-terminal 5 amino acid “message” and generate numerous product peptides that have been shown to strongly antagonize signaling through the FLS2-BAK1 co-receptor ^13,14,61^. Flg22 disarmament by IssA at least partially explains the ability of *Dja* MF79 to suppress flg22-induced plant immunity and promote root colonization by other microbes ^21^. In a similar manner, the plant pathogen *Pseudomonas syringae* pv. *tomato* DC3000 utilizes the Type I secreted zinc metalloprotease effector AprA to degrade flagellin monomers and blocks flagellin-induced MTI ^62^. *Dja* MF79, however, has not been observed to cause disease and is a strong endophytic colonizer of Arabidopsis ^21,42-44^. Flg22 from *Dja* MF79 has an altered “message” peptide GLAIS relative to the canonical *P. aeruginosa* flg22 sequence, but is still mildly immunogenic ^9^ and would presumably be cleaved by IssA. However, *issA* deletion did not cause *Dja* MF79 to either activate plant immunity or cause root growth inhibition (Figure 3A, Figure 4B, Figure S2B).

IssA-like subtilases are found in the majority of suppressive isolates found in our 165-member collection, especially in isolates in order Xanthomonadales (Figure 4A, Table S4-S5). We also found homologs of IssA in the majority of previously reported suppressive isolates from order Xanthomonadales (Table S4) ^22^. IssA-like subtilases are not limited to order Xanthomonadales, as many other isolates in the 165-member collection contain at least one homolog (Table S5). Deletion of *issA* in *Dja* MF79 and IssA-like subtilases from genetically tractable *Luteibacter* sp. MF22 (a Xanthomonadales order of Gammaproteobacterium) and *Ralstonia* sp. CL21 (a Betaproteobacterium) resulted in a reduction of suppression of flg22-induced *pCYP71A12::GUS* activation (Figure 4B). While IssA homologs are present in many suppressors identified in our screen, there are notable exceptions. The three strongly suppressive *Curtobacterium* spp. CL17, CL20, and MF314 (Actinobacteria, order Micrococcales) lack any homolog of IssA, suggesting a separate mechanism of flg22-induced MTI suppression.

Our results suggest a model where IssA-like subtilases are secreted by various commensal bacteria to rapidly cleave flagellin prior to perception by the plant PRRs and activation of immunity. This represents a new mechanism of innate immune suppression by commensal microbes and adds support to the theory that suppressive commensals like *Dja* MF79 act as negative regulators of plant immunity, enabling plants to balance growth and defense ^21,22^. In a dysbiotic microbiome caused by an increase in pathogen abundance, the relative decrease of commensal immunosuppressors could lead to a corresponding increase in flg22-induced immune activation. In this way, immunosuppressive commensal rhizobacteria like *Dja* MF79 might help shape plant receptor-mediated decision making by self-regulating immunogenic MAMPs, allowing the host to engage with beneficial and nonharmful bacteria while simultaneously restricting harmful bacteria ^4,6^.

## METHODS

### Bacterial strains and culture conditions

Bacterial strains were routinely cultured on 0.5× TSB medium (15 g/L tryptic soy broth CM0129B w/ or w/o 20 g/L agar) or LB medium (10 g/L tryptone, 5 g/L yeast extract, 5 g/L sodium chloride w/ or w/o 15 g/L agar) at 28 °C. *E. coli* strains were routinely cultured on LB media at 37 °C. *E. coli* strain WM3064 was cultured on LB medium including 300 µM diaminopimelic acid (DAP). The following antibiotics were included when appropriate at the following concentrations: Kanamycin (Km) 50 µg/mL and Chloramphenicol (Cm) 33 µg/mL.

### Plant growth conditions

Plants were grown in growth chambers either under short-day conditions with a 10-hour light/14-hour dark, 21 °C/18 °C day/night cycle or under long-day conditions with a 16-hour light/8-hour dark, 21 °C/18 °C day/night cycle. Light intensity was 170 μmol·m^-2^·s^-1^ and relative humidity was kept at 70%.

### Root growth inhibition assay

Seeds of an *A. thaliana* line expressing a *pWER::FLS2-GFP* construct ^46^ were surface-sterilized for 10 minutes with a solution of 70% bleach and 0.2% tween-20. Seeds were washed five times with sterile ddH_2_O and germinated on 0.5× MS media (2.22 g/L PhytoTech Labs M404 Murashige & Skoog modified basal medium with Gamborg vitamins, 0.5 g/L MES, 10 g/L agar, 5 g/L sucrose, (pH 5.6-5.7)) in 12×12 cm square petri plates in vertical orientation under short-day conditions for 7 days. 165 root-associated bacterial isolates (Table S1) ^2^ were streaked from 20% glycerol stocks onto agar plates and incubated at 28 °C for 2-4 days. A single colony was picked and inoculated in 5 ml of liquid medium and grown at 28 °C and 250 rpm for two or three days. Bacterial cultures were pelleted by centrifugation at 5,000× G for 10 min and washed two times in 3 ml of 10 mM sterile MgCl_2_. The OD_600_ was measured by nanodrop and normalized to OD_600_=0.01 in 10 mM sterile MgCl_2_. 100 µL of bacteria was spread on four 0.5× MS plates (without sucrose) per strain, with half of the plates containing 1 μM flg22 and the other half containing no flg22. 8 to 10 germinated Arabidopsis *pWER::FLS2-GFP* seedlings were transferred from germination plates to bacteria-inoculated plates and the initial root tip location was marked. The plates containing the seedlings and bacteria were sealed with 3M tape (Thermo Fisher) and grown in vertical orientation under short-day conditions for seven days. The plates were imaged using a camera and the length of primary root was measured using ImageJ by freehand lines to quantify the change in root tip position from the initial to the final location (Table S2).

### pCYP71A12::GUS assay

GUS assays were conducted as described by Teixeira et al. ^21^. Seeds of an *A. thaliana* line carrying a *pCYP71A12::GUS* reporter construct ^47^ were surface-sterilized for 10 minutes with a solution of 20% bleach and 0.5% tween-20. Seeds were washed three times with sterile ddH_2_O, then two to three seeds were added to wells of a 24-well microplate and grown in 500 µL of seedling growth media (SGM) (4.44 g/L MS salt, 0.5 g/L MES, 5 g/L sucrose, pH adjusted to 5.7 with KOH). Plants were grown under long-day conditions for 6 days. In assays using bacteria culture, the SGM was replaced on the sixth day with fresh SGM and bacteria that had been grown overnight at 28° in LB media, washed three times with SGM, and adjusted to the appropriate density. Inoculated plants were incubated overnight in long-day conditions prior to flg22 induction. In assays using cell-free bacterial culture supernatant, the supernatant was prepared as described below and added on the sixth day concurrently with fresh SGM and flg22. In assays using purified, recombinant IssA, the protein was added concurrently with fresh SGM and flg22. In all experiments, plants were incubated with flg22 for 5 hours under long-day conditions, after which they were washed with 1000 µL of 50 mM sodium phosphate buffer (pH 7.0) and incubated in 500 µL of GUS buffer (50 mM sodium phosphate (pH 7.0); 10mM EDTA; 0.5 mM K_4_ ferricyanide; 0.5 mM K_3_ ferricyanide; 500 µM X-gluc in DMF; and 0.01% Silwet L-77). Plants were incubated overnight in the dark at 37 °C. The following day the GUS buffer was replaced with 500 µL 3:1 ethanol: acetic acid and left at 4 °C overnight. The plants were then imaged in 95% ethanol using an Amscope FMA050 dissecting microscope.

### Bacterial mutagenesis

Gene deletion and knock-in was conducted as described by Colaianni et al.^9^. DNA 1 kb -upstream and -downstream from *issA* and *issA2* were PCR-amplified from *Dja* MF79, *Luteibacter* sp. MF22, and *Ralstonia* sp. CL21 using Q5 Hi Fidelity DNA Polymerase and cloned into the SacB-containing vector pMO130 ^63^by Gibson assembly with NEB HiFi DNA assembly. The vector for deletion of *issA* from *Dja* MF79 additionally included a chloramphenicol resistance cassette. The resulting vectors were transformed into *E. coli* WM3064, conjugated into host strains using bi-parental mating, and integrated into the host chromosomes by homologous recombination. Resulting colonies were re-plated on LB medium containing 10 g/L sucrose and 0.1 mM IPTG to counter-select for vector excision. The final mutants were confirmed by PCR.

### Expression and purification of recombinant IssA

Construction of a heterologous *P. pastoris* IssA expression strain and secreted expression of recombinant IssA were as described previously for fungal subtilase Vd-DUMP, except that protein expression was carried out for 4 days at 15 °C ^56^. Briefly, a synthetic DNA was cloned into pPICZaA expression plasmid to create pTAN282. Linearized pTAN282 was transformed into *P. pastoris* X-33 followed by selection on zeocin-containing plates. Cells from eight colony-forming units were tested for IssA expression. All strains secreted similar amounts of IssA, the best was retained as TAN670.

Following expression from TAN670, IssA was precipitated from 1 L cell-free media by salting out with ammonium sulfate (351 g/L). Precipitated protein was dissolved in 20 ml phosphate buffered saline and dialyzed (16 h; 4°C). After dialysis, IssA was further purified by precipitation with cold acetone (1.5 V; -20°C). Precipitated protein was dissolved in 20 ml anion exchange buffer (20 mM Tris-Cl, pH 8.0) and loaded onto a HiTrap Capto Q (1 ml) anion exchange column. The majority of IssA did not bind, while media components were removed. The flow through containing IssA was used in bioassays and enzyme assays. IssA concentrations were determined by measuring Abs_280_, with an extinction coefficient of 63,620 based on the amino acid sequence ^64^. Aliquots were stored at -80°C. We did not observe loss of IssA activity after multiple freeze/thaw cycles.

### Protein LC/MS/MS

In-gel digestion of protein bands using trypsin was performed as in Shevchenko et al. ^65^. Trypsin digested samples were dried completely in a SpeedVac, resuspended with 20 µl of 0.1% formic acid pH 3 in water, and 2 µL (∼ 360ng) was injected per run using an Easy-nLC 1200 UPLC system. Samples were loaded directly onto a 45cm long 100um inner diameter nano capillary column packed with 1.9um C18-AQ resin (Dr. Maisch, Germany) mated to metal emitter in-line with an Orbitrap Fusion Lumos (Thermo Scientific, USA). The column temperature was set at 45 °C with a 1-hour gradient method and 360 nL/min flow rate. The mass spectrometer was operated in data dependent mode with the 120,000 resolution MS1 scan (positive mode, profile data type, AGC gain of 4e5, maximum injection time of 54 sec and mass range of 375-1500 m/z) in the Orbitrap followed by HCD fragmentation in ion trap with 35% collision energy. A dynamic exclusion list was invoked to exclude previously sequenced peptides for 60s and maximum cycle time of 3s was used. Peptides were isolated for fragmentation using quadrupole (1.2 m/z isolation window). Ion-trap was operated in Rapid mode. Raw files were searched using Sequest-HT algorithms ^66^ within the Proteome Discoverer 2.5.0 suite (Thermo Scientific, USA). 10 ppm MS1 and 0.6 Da MS2 mass tolerances were specified. Carbamidomethylation of cysteine was used as fixed modification, oxidation of methionine, deamidation of asparagine and glutamine were specified as dynamic modifications. Pyro glutamate conversion from glutamic acid and glutamine were set as dynamic modifications at peptide N-terminus. Acetylation was specified as dynamic modification at protein N-terminus. Trypsin digestion with maximum of 2 missed cleavages were allowed. Files were searched against *E. coli* (K12) database downloaded from Uniprot.org with both the full-length IssA and IssA beginning at T157 protein sequences added.

### Casein clearance assay

Bacteria strains were grown in LB liquid cultures overnight and inoculated at OD_600_=1.0 onto plates containing 10 g/L Casein, 2.5 g/L yeast extract, 5 g/L tryptone, 10 g/L glucose, 10 g/L agar, (pH 7.0) with a hole punched out of the center using a sterile cork borer. Bacteria were incubated in the plates for 7 days at 28 °C, after which plates were washed with ddH_2_O and imaged with overhead white LED lights in a Syngene GBox. The radius of clearance was measured using ImageJ.

### Preparation of cell-free bacterial supernatant

Bacteria strains were grown in LB liquid cultures overnight, washed three times with 10 mM MgCl_2_, and inoculated at OD_600_=0.02 into SGM media containing 0.1% tryptone. Strains were grown at 28 °C and 250 rpm for exactly 18 hours. The bacteria were centrifuged for 15 minutes at 4000 g and the supernatant was filtered with a 0.22 µm filter.

### Flg22 supernatant degradation assays

Cell-free bacterial supernatant was collected as described above. After growing plants for 6 days as described above in the GUS assay method, 0.1% supernatant in 600 µL ice-cold SGM was mixed with 100 nM flg22 in a Pierce 10 kDa MWCO protein concentrator (Thermo Scientific). The mixture was incubated for 30 seconds than passed through the concentrator at 5000×G. 500 µL of the permeate was added to 6-day old *pCYP71A12*::*GUS* seedlings and the GUS histochemical assay was conducted as described above.

### MALDI-TOF MS-based protease assays

Protease reactions contained flg22 (1 mM) in reaction buffer (10 mM sodium acetate, pH 5.2). Reactions were initiated by addition of IssA (1 µM). Time points were collected from reactions by removing 10 µL aliquots and combining with an equal volume of matrix (saturated 2,5-dihydroxybenzoic acid in acetonitrile). Time points were spotted onto a target, dried, and analyzed using a Bruker-Daltonics Microflex LRF with a pulsed N2 laser with reflectron acquisition (3000 shots, 337 Hz, 60 Hz pulse).

## ACKNOWLEDGEMENTS

We are grateful to Jeff Dangl (University of North Carolina Chapel Hill) for the strains in the 165-member isolate collection used in this study. We thank Saw Kyin in Princeton’s University Proteomics and Mass Spectrometry Core Facility for performing protein LC/MS/MS analysis and Kurt Sollenberger for technical assistance with recombinant protein expression and purification.

This work was supported in part by Princeton University startup funding to J.M.C. This work was supported in part by the U.S. Department of Agriculture, Agricultural Research Service.

K.F., B.J., and I.Y. acknowledge support from the Princeton University Lindow Senior Thesis Fund.

K.F. acknowledges support of the High Meadows Environmental Institute at Princeton University.

## AUTHOR CONTRIBUTIONS

Conceptualization: S.E., T.J., and J.M.C.; Methodology: S.E., T.J., K.F., T.N., and J.M.C.; Validation: K.F., Investigation: S.E., T.J., K.F., C.L., S.W., Y.O.B., A.E., B.J., I.Y., and T.N.; Writing – Original Draft: S.E.; Writing – Review and Editing: S.E., T.J., K.F., T.N. and J.M.C.; Supervision: J.M.C.

