## Supplementary Figures for "A type II secreted subtilase from commensal rhizobacteria disarms the immune elicitor peptide flg22"

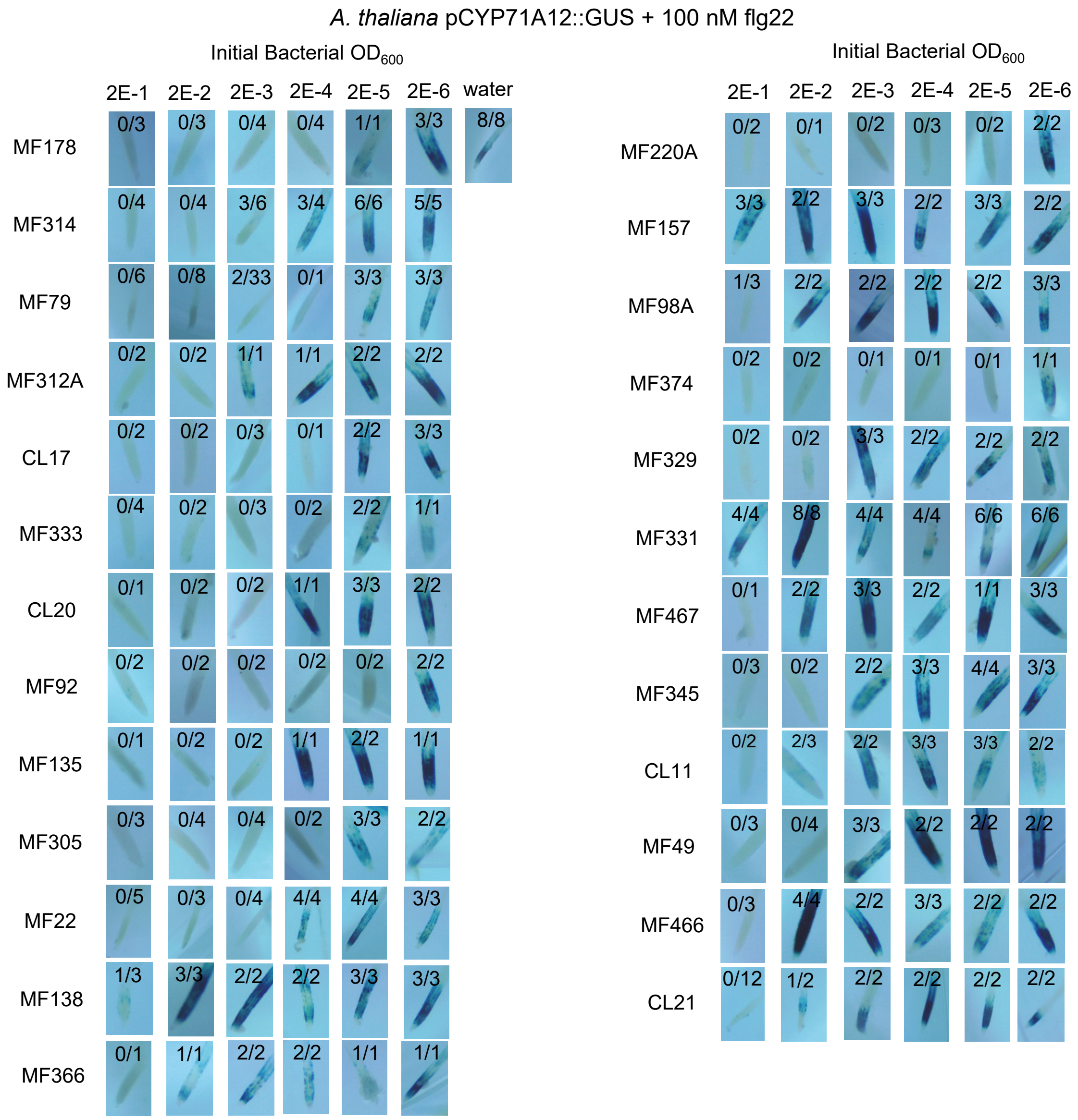


**Figure S1: Representative images of Arabidopsis *pCYP71A12::GUS* roots from Figure 2B.** Plants were germinated in sterile growth liquid media for six days, then the growth media was replaced with media containing bacteria at the indicated densities. The next day, plants were induced with 100 nM flg22 for five hours then incubated with GUS reagent overnight. Plants were destained and root tips were photographed under a microscope. Numbers indicate the number of roots displaying visible GUS expression.

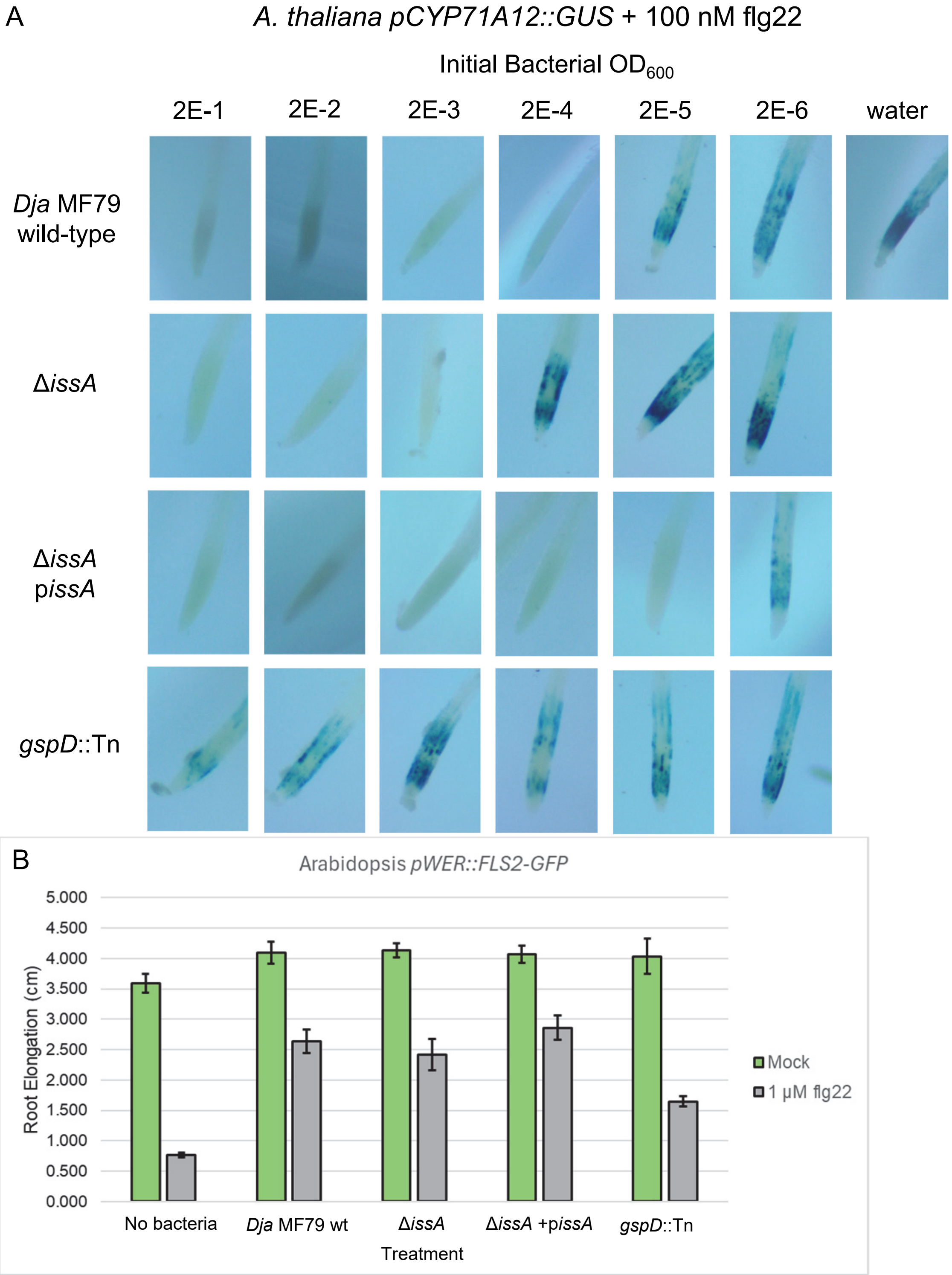


**Figure S2: *issA* contributes to *Dja* MF79 suppression of flg22-induced *pCYP71A12::GUS* activation but not flg22-induced RGI.** (A) GUS assay utilizing Arabidopsis *pCYP71A12:GUS* roots and *Dyella japonica* strains. *issA* deletion increased the inoculation density of bacteria need to suppress flg22-induced *pCYP71A12*::*GUS* activation by an order of magnitude. Complementation with p*issA* decreased the inoculation density of bacteria needed to suppress the responses by two orders of magnitude. *gspD::Tn* mutants were unable to suppress responses at all bacteria densities tested. (B) RGI assay utilizing Arabidopsis *pWER:FLS2-GFP* plants and *Dyella japonica* strains. Wild-type *Dja* suppressed root growth inhibition by 1 µM flg22. Δ*issA* strains did not have observably different RGI suppression than wild-type. Errors bars indicate standard error. The experiment was repeated three times with similar results.

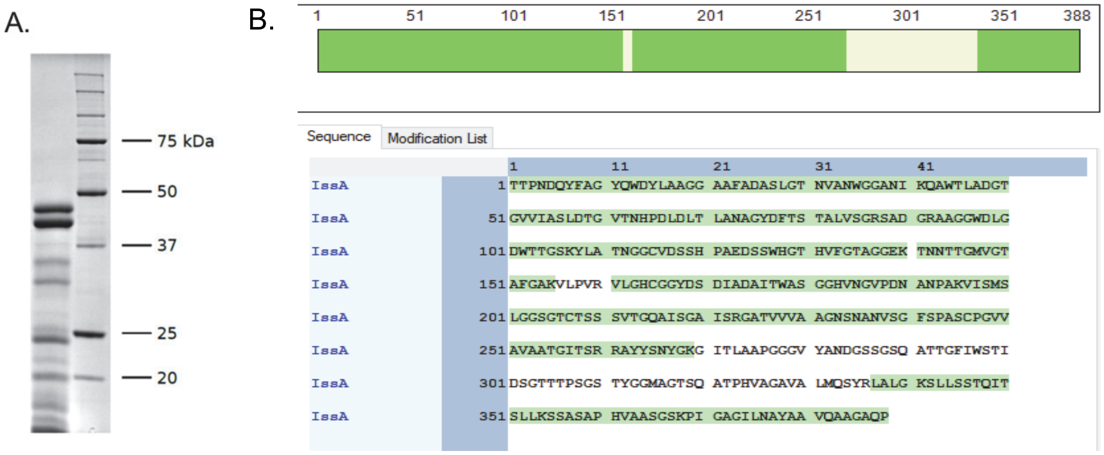


**Figure S3. Purified recombinant IssA exhibits autocleavage activity.** (A) Gel electrophoresis of purified, recombinant IssA from *Pichia pastoris*. (B) Coverage map of the largest two bands from (A). Recombinant IssA was purified from *Pichia pastoris* and run on an SDS-PAGE gel. Coomassie-stained bands were assessed by LC-MS timsTOF. Coverage of peptides is indicated in green. Both of the largest bands in (A) produced identical coverage maps.

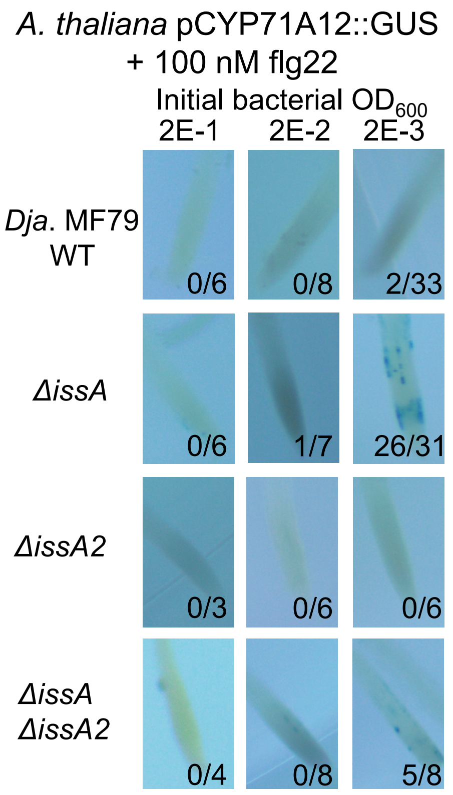


**Figure S4: Representative images of Arabidopsis *pCYP71A12::GUS* roots from Figure 4.** Plants were germinated in sterile growth liquid media for six days, then the growth media was replaced with media containing bacteria at the indicated densities. The next day, plants were induced with 100 nM flg22 for five hours then incubated with GUS reagent overnight. Plants were destained and root tips were photographed under a microscope. Numbers indicate the number of roots displaying visible GUS expression.

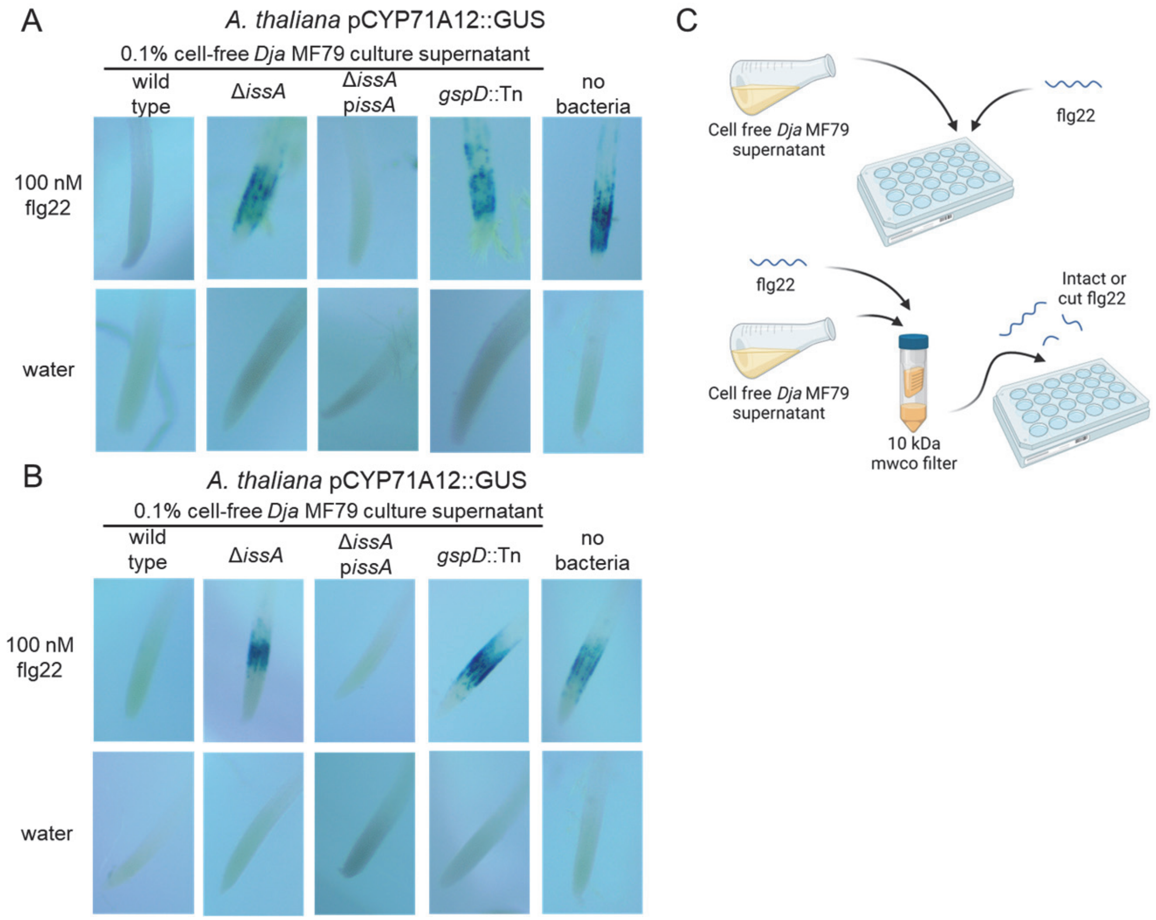


**Figure S5: *Dja* MF79 suppresses flg22 immunogenicity *in vitro* in a IssA-dependent manner**. (A) GUS assay utilizing cell-free culture supernatant. Arabidopsis *pCYP71A12:GUS* plants were grown for six days in sterile growth liquid, then the growth media was replaced with media amended with 100 nM flg22 or water and 0.1% overnight bacterial culture supernatant filtered with a .22 µm filter. Cells were induced for five hours then incubated overnight with GUS reagent, destained, and imaged. (B) GUS assay utilizing flg22 co-incubated with cell-free culture supernatant. 0.1% cell-free bacterial culture supernatant was incubated with 100 nM flg22 or water for one minute, then the mixtures were applied to a 10 kDa MWCO filter. The filtrate was incubated with six-day old Arabidopsis *pCYP71A12:GUS* plants for five hours, incubated overnight with GUS reagent, destained, and imaged. (C) Cartoon of methods used in (A) and (B). Image created with BioRender.com
